# Singing above the chorus: cooperative Princess cichlid fish (*Neolamprologus pulcher*) has high pitch

**DOI:** 10.1101/039313

**Authors:** Rachel K. Spinks, Moritz Muschick, Walter Salzburger, Hugo F. Gante

## Abstract

Teleost fishes not only communicate with well-known visual cues, but also olfactory and acoustic signals. Communicating with sound has advantages, as signals propagate fast, omnidirectionally, around obstacles, and over long distances. Heterogeneous environments might favour multimodal communication, especially in socially complex species, as combination of modalities’ strengths helps overcome their individual limitations. Cichlid fishes are known to be vocal, but a recent report suggests that this is not the case for the socially complex Princess cichlid *Neolamprologus pulcher* from Lake Tanganyika. Here we further investigated acoustic communication in this species. Wild and captive *N. pulcher* produced high frequency sounds (mean: 12 kHz), when stimulated by mirror images. In laboratory experiments, *N. pulcher* produced distinct two-pulsed calls mostly, but not exclusively, associated with agonistic displays. Our results suggest that male *N. pulcher* produce more sounds at greater durations than females. Thus, we confirm that the Princess cichlid does not produce low frequency sounds, but does produce high frequency sounds, both in combination with and independent from visual displays, suggesting that sounds are not a by-product of displays. Further studies on the hearing abilities of *N. pulcher* are needed to clarify if the high-frequency sounds are used in intra-or inter-specific communication.

## Introduction

In spite of the long-held view of a silent underwater world, we now know that many teleost fishes produce sounds as part of their normal behavioural repertoire (Lobel et al., 2010). It should come as no surprise that fish ubiquitously use sounds to communicate, as water is a superior acoustic medium, where sound travels almost five times faster than in air (Fine & Parmentier, 2015). Compared to other signal modalities auditory signals can present some advantages: they propagate fast and in all directions unlike olfactory cues, in which case the receiver must be downstream from the sender (Fine & Parmentier, 2015); or around obstacles and to longer distances than visual signals, which quickly become attenuated with increasing distance, in low light or in deep water conditions (Lythgoe & Partridge, 1991). For instance, the nocturnal New Zealand bigeye fish (*Pempheris adspersa*) produces sounds mainly at night to promote shoal cohesion when visual cues have reduced utility (Radford et al., 2015).

Nevertheless, long-range auditory signals also present some communicative weaknesses. For instance, fish need to deal with high levels of environmental noise in shallow water habitats (Ladich & Schulz-Mirbach, 2013; Lugli, 2015) and there is the potential for eavesdropping by non-intended receivers, conspecifics or predators (Verzijden et al., 2010; Bradbury & Vehrencamp, 2011; Maruska et al., 2012). The alternate or simultaneous use of signals of different modalities combines their strengths and reduces limitations imposed by the environment on a particular type of signal (Stevens, 2013). Multimodal communication is thus expected to evolve under varied and unstable environments (Munoz & Blumstein, 2012), in particular in gregarious, territorial and socially complex species (Freeberg et al., 2012).

Fish commonly produce sounds in agonistic, reproductive and defensive contexts (Lobel et al., 2010), either in isolation or most often in association with visual signals (Ladich, 1990, 1997). Such sounds are usually low frequency purrs and grunts (40-1000 Hz), but higher frequency clicks and creaks (above 1 kHz) have also been reported (Ladich, 1997; Lobel et al., 2010). A group of fish that has received increasing attention regarding sound production are cichlids. In particular, those originating from the East African Great Lakes are prime models for studying diversification and adaptation due to varied life histories, morphologies and behaviours (Salzburger, 2009; Gante & Salzburger, 2012). While diversity in colour patterns and visual adaptations have long been recognised as a driving force in cichlid evolution (Santos & Salzburger, 2012; Wagner et al., 2012), the description of sound production and hearing abilities have only more recently gained momentum in spite of a long history of research (Amorim, 2006; Ladich & Fay, 2013).

Here we report on the production of sounds by the Princess cichlid, *Neolamprologus pulcher* (Trewavas & Poll, 1952). This cooperatively breeding species lives in rocky shores of southern Lake Tanganyika, East Africa, home to one of the most diverse freshwater fish adaptive radiations (Muschick et al., 2012; Salzburger et al., 2014), and has become a favourite in studies of animal cooperation (Wong & Balshine, 2011; Zöttl et al., 2013). In *N. pulcher* each extended family is typically formed by a dominant breeding couple and up to a few dozen subordinate helpers that collectively raise young and defend their territory from other such groups in the colony. Considering the heterogeneous nature of rocky habitats (especially when compared to sandy habitats) and the high social complexity of cooperative breeders, *N. pulcher* is expected to show increased levels of communicative complexity. Indeed it has been shown that Princess cichlids use a combination of visual and olfactory signals or cues in multiple aspects of their lives, such as individual recognition, territoriality and aggression (Bachmann et al., (n.d.); Balshine-Earn & Lotem, 1998; Frostman & Sherman, 2004; Le Vin et al., 2010). It is thus puzzling that *N. pulcher* have reportedly gone completely silent (Pisanski et al., 2014). In this study we further investigate the possibility of acoustic communication in this species by examining both captive-bred and wild-caught fish, over a much wider range of sound frequencies than before.

## Methods

### Acoustic recordings of wild-caught *N. pulcher* - field experiments

Recordings of wild *N. pulcher* were conducted in July and August 2013. *Neolamprologus pulcher* from different social groups were carefully captured with gill nets on SCUBA in shallow waters around Kalambo Lodge, Isanga Bay, Zambia, in the south-eastern shore of Lake Tanganyika (8°37’22.1”S, 31°12’03.6”E). Around 20 adult fish were placed together in a concrete pond (1 × 1 × 1 m), with lake water and without shelters, so aggression levels were reduced between individuals, and left to acclimatise for 3 days before the recordings commenced. Every second day, one-third of the water in the pond was changed. Fish were individually recorded in another concrete pond that was the same size, but only filled to 20 cm depth. An octagonal arena, with mirror panels (25 × 20 cm) on the inside, was used to elicit behaviours and sounds (Fig. 1A). Mirrors have been successfully used to induce typical agonistic behaviours in African cichlids and fish in general (Rowland, 1999; Dijkstra et al., 2012). Contrary to the use of interacting, live fish as stimuli, mirrors have the advantage that sound emitters can not be mistaken, and because only one individual is recorded at any one time, precise calculation of sound parameters is also facilitated. To prevent the fish from seeing multiple mirror images, a perforated box was placed in the centre of the arena (Fig. 1A). A Teledyne Reson TC4013 hydrophone (Denmark), with a receiving sensitivity of −211 dB re: V/μPa and frequency range of 1 Hz to 170 kHz, was suspended inside the perforated box. Sound was intensified at 500 Hz by an UltraSoundGate charge amplifier and then stored and digitalised at 48 kHz (with 16 bit resolution) into Waveform Audio File Format (.wav) by the Marantz PMD670 recorder. Movements were recorded from above with a GoPro Hero 3 camera that was synchronised to the sound recordings. This allowed discarding sounds that had been produced by the fish touching the setup or breaking the water surface. The pond was illuminated with indirect natural daylight and two solar-charged LED lamps. Unlike fluorescent bulbs, LEDs produce negligible levels of low frequency sound (Rumyantsev et al., 2005).

**Fig. 1.**
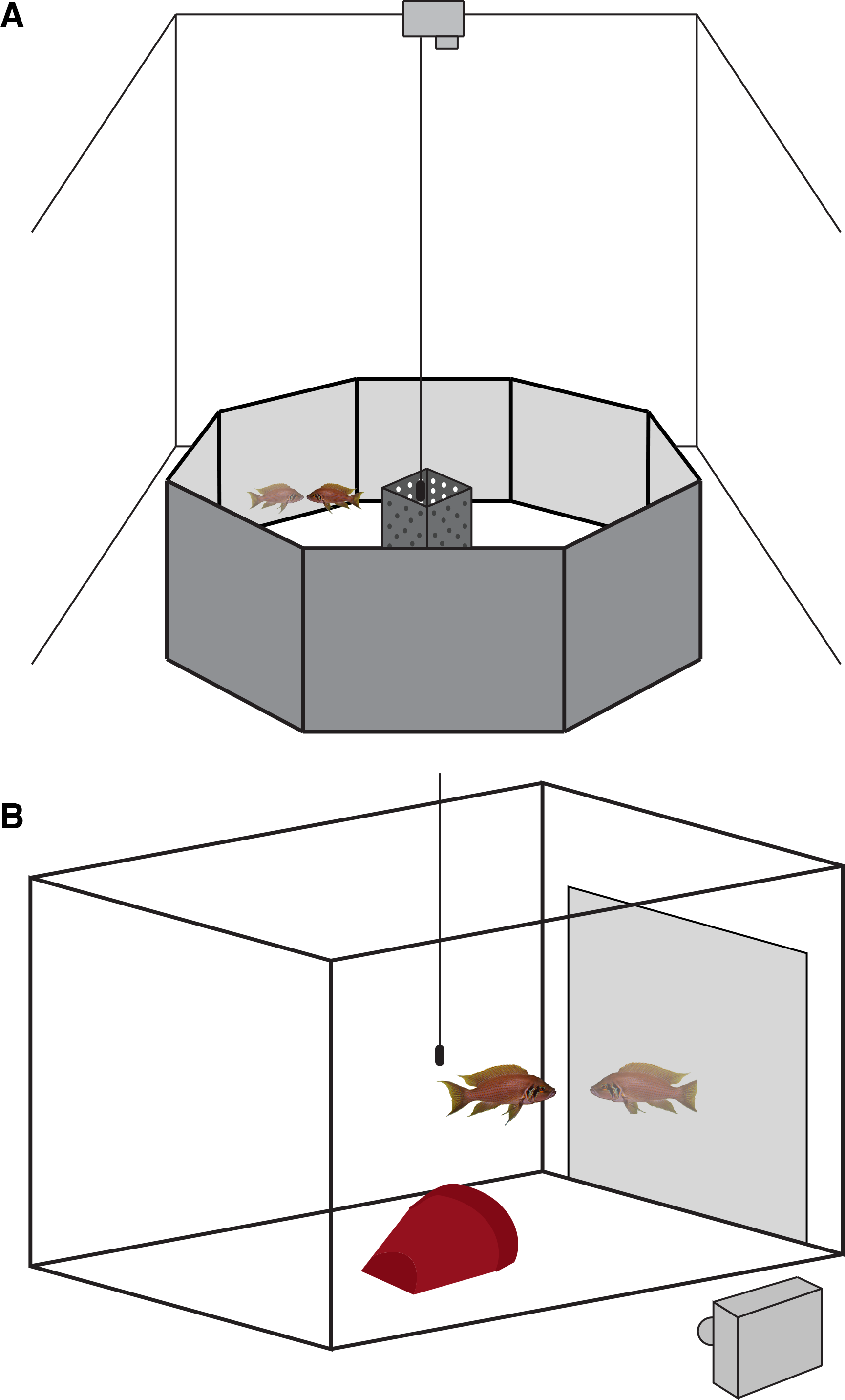
Setups used to record sounds produced by *N. pulcher.* In the field experiment (A), an octagonal mirror arena was used, while in the laboratory experiment (B), one glass mirror was placed against a wall of the aquarium

Individuals were introduced to the experimental arena via a box with a sliding door. After a 2-minute acclimatisation period the door of the box was opened and the box removed as soon as the fish had vacated it. If the fish did not exit right away, the box was lifted slightly to encourage departure. Each fish was recorded for eight minutes and then weighed, standard length measured, and sexed by examining the genital papilla. A total of ten (6 males and 4 females) *N. pulcher* were used in this study. Recordings of wild fish taken at Lake Tanganyika were first manually inspected for sounds and then filtered with a bandpass at 300 Hz to remove low frequency background noise. The experiments were done in accordance with the Department of Fisheries, Lake Tanganyika Research Unit, Mpulungu, Zambia.

### Acoustic recordings of captive-raised *N. pulcher*- laboratory experiments

Given the recent report of silent *N. pulcher* (Pisanski et al., 2014), sound recordings were repeated under laboratory conditions, where a camera could be placed in lateral view to monitor fish behaviours with greater detail than in the field (Fig. 1B). It also allowed controlling for the effect of captive raising on sound production.

In order to minimise ambient background noise, acoustic recordings took place in a room with thick concrete walls, with an aquarium (40 × 30 × 25 cm) resting on 2 cm-thick acoustic absorption cotton and placed inside a large (48 × 42 × 32 cm) expanded polystyrene foam box. The inside of the expanded polystyrene foam container, except for the floor, was also covered with acoustic insulation that allowed external sounds to be reflected and internal sounds to be absorbed to reduce reverberation. Four battery-operated LED lamps were placed above the aquarium to provide adequate illumination. The aquarium contained a half terracotta flowerpot to provide shelter for the fish.

First or second generation laboratory-raised *N. pulcher* were used, originating from fish collected at Kalambo Lodge, Isanga Bay, Zambia in Lake Tanganyika. Fish were originally kept in pairs in aquaria with sandy substrate, halved terracotta flowerpots and a motorised sponge filter, and were fed once daily prior to the experiment. Ten sexually mature *N. pulcher* (5 males and 5 females) were then selected and individually recorded in April 2015. A 1.9 mm-thick glass mirror (28 × 22 cm), placed flat against a lateral wall inside the aquarium, was used to induce sound production (Fig. 1B). Fish were gently hand-netted from their home aquaria and given one hour to acclimatise in the experimental setup; however, the mirror was introduced to the aquarium only two minutes before the recording began to prevent the fish becoming accustomed to it. All nearby electrical equipment, including the room lights, were shut off shortly before synchronous video and audio recordings commenced.

We used the same hydrophone, amplifier, recorder and settings as described in the field experiment. Although in the laboratory recordings we utilised the Raven Pro 1.5 sound analysis software’s adaptive broadband filter, with the default settings of a filter order of ten and a least mean squares step size of 0.01, to reduce the likelihood of filtering out potential fish sounds (Bioacoustics Research Program, 2014). Adaptive broadband filtering is useful when the preferred broadband signal is amidst narrowband background noise that could not otherwise be eliminated (Bioacoustics Research Program, 2014). This filter works just like when people talk in a noisy environment, the continuous surrounding background sounds are recognised but the focus and concentration is on the person’s speech, or in this case on the sounds produced by the fish. To diminish distortion of the fish’s acoustic signals in the aquarium, the hydrophone was placed within the attenuation distance of where the fish were expected to produce sound (Akamatsu et al., 2002). Behaviour was simultaneously recorded with a Nikon 1 camera with an 11-27.5 mm lens. Each recording session lasted 20 minutes. Subsequently, fish were weighed, standard length measured, sexed by examination of the external genital papilla and then returned to their home aquarium. Experiments were authorised by the Cantonal Veterinary Office, Basel, Switzerland (permit numbers 2317 & 2356).

### Characterisation of *N. pulcher* sounds

Only sounds that showed a clear structure and high signal to noise ratio were considered. All sounds were confirmed with the synchronised video footage and if, for example, the fish touched the mirror or turned around quickly, resulting in an incidental sound, or an unexpected background noise occurred, then no measurements were taken at this time. For this reason, we focused on characterising sounds produced by fish only in the laboratory experiment, where behaviours could be unequivocally monitored. Based on the typical social behaviours of *N. pulcher* (Table 1) we noted if a behavioural display was associated with sound. To quantify the acoustic properties of sounds produced by *N. pulcher* in the laboratory we measured pulse duration, pulse peak frequency, interpulse interval, call duration, and pulse rate (Fig. 2). In the field dataset, we focused on pulse duration and pulse peak frequency. In our study the duration of each pulse is defined as the time in milliseconds (ms) from the onset of a pulse to its end as classified by amplitude of the signal. Pulse peak frequency is the frequency with the maximum power in the pulse. The duration between each pulse, the interpulse interval, is calculated in milliseconds and is the period with only white noise levels of sound between the pulses. The duration of a call, in milliseconds, is measured from the onset of the first pulse to the end of the last pulse and may contain one pulse or many. Call duration is often subjectively measured in fish acoustics literature. We aimed to provide a non-biased, replicable classification by measuring every interpulse interval in the recordings (these periods of white noise went from milliseconds to minutes) and plotting their frequencies as a histogram. Any discontinuity would be indicative of how many pulses constitute a typical call. Lastly the pulse rate can be defined as the function of the number of pulses per call duration.

**Table 1.**
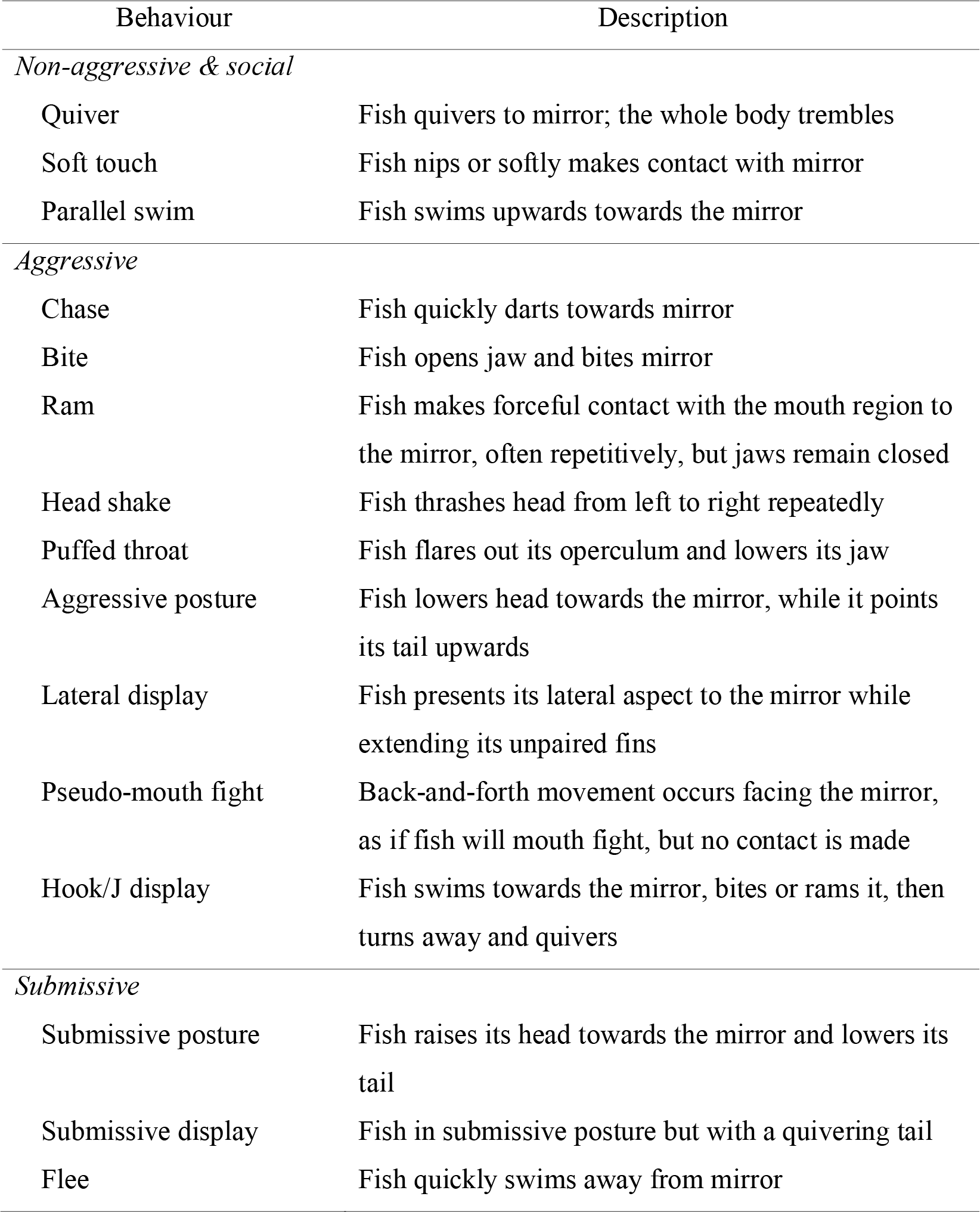
*Neolamprologus pulcher* ethogram illustrates typical social behaviours of the species (adapted from (Sopinka et al., 2009; Pisanski et al., 2014))

**Fig. 2.**
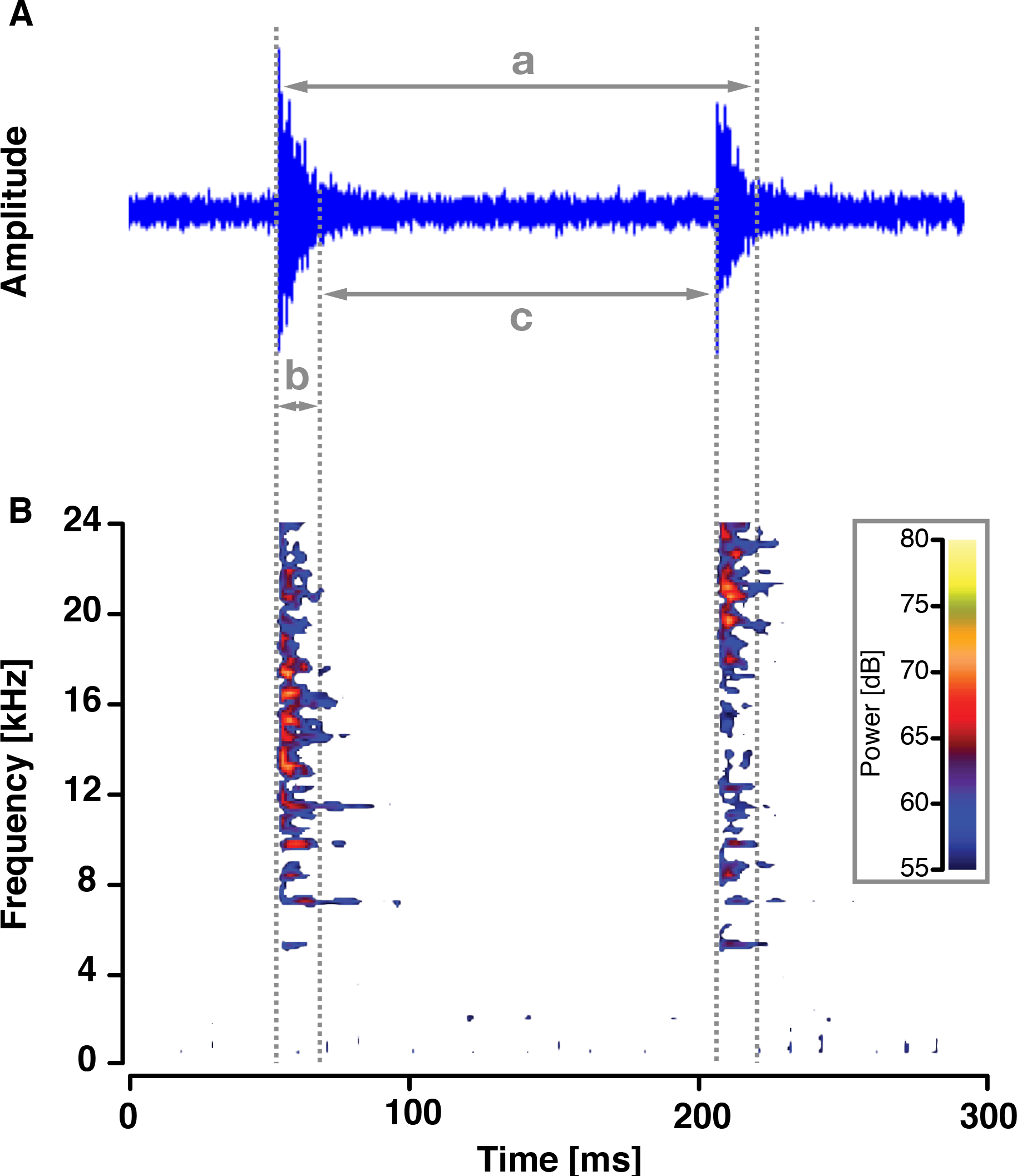
Oscillogram and spectrogram of a sound produced by *N. pulcher.* The oscillogram (A) presents the waveform of the pulses in time versus amplitude. Whereas the spectrogram (B) shows how the frequency of the pulses changes over time, and the colour indicates the relative amplitude. Here, the aforementioned temporal parameters; call duration (a), pulse duration (b) and interpulse interval (c) are illustratively defined. This double-pulsed call was made by a male in the laboratory experiments that concurrently exhibited an aggressive lateral display just after a series of rams and bites to the mirror

The aforementioned temporal parameters were measured on the oscillogram in the same preset window size and settings. Whereas peak frequency was quantified with the spectrogram (Hann, FFT size 256 samples, filter bandwidth 270Hz, with a 50% overlap). All measurements were made in Raven Pro 1.5 sound analysis software, commonly employed in animal communication research (Bioacoustics Research Program, 2014).

### Results

Of the seven (four males and three females) out of 10 *N. pulcher* that produced sound in our setup at Lake Tanganyika, there were a total of 40 pulses recorded (mean ± SD; 5.7 ± 7.1 pulses/fish). Mean pulse duration was 1.5 ± 0.5 ms, whilst pulse peak frequency was 12008.0 ± 8312.8 Hz. In the laboratory setting, six (four males and two females) out of 10 *N. pulcher* emitted sound. Of those six individuals, five produced sound associated with a defined social behaviour (Table 2). Sound production occurred most frequently when fish were in an aggressive posture or lateral display (Table 2). Often, these aggressive displays coupled with sound production were followed by or occurred shortly before other aggressive behaviours such as rams, bites and chases. Males only exhibited aggressive behaviours coupled with sound, whereas females in addition showed submissive displays in conjunction with sound (Table 2). One female predominantly produced sound alongside non-aggressive social and submissive behaviours (Table 2). Five doubled-pulsed calls from two fish (one male and one female) were also recorded without concurrent visual display, when both fish were motionless (Table 3). This particular female had produced sound with behavioural displays, however paused displaying for a couple of minutes and continued to call and then began displaying again. The male on the other hand did not display once, he performed a few exploratory swims of the aquarium and then stayed in the corner of the aquarium calling out the rest of the recording. These sounds did not come from background or incidental noise and were similar to the other acoustic signals produced during displays (Table 3).

A total of 92 pulses (14.8 ± 11.5 pulses/fish) produced by six individuals were measured in the laboratory setup (Table 3). Since the minimum resonance frequency of the aquarium (~4000 Hz) was much lower than the dominant frequency of *N. pulcher* sounds (~12000 Hz, Table 3), according to (Akamatsu et al., 2002) resonance distortion in the aquarium should be minimal. Inspection of interpulse duration frequency revealed that the majority of pulses were produced less than 0.4 s apart (Fig. 3). Pulses separated by less than 0.4 s were then considered part of one call, and on average 2 pulses were produced per call (Additional File 1). When this double-pulse call occurred, often the first pulse had a dominant frequency between 7000 Hz and 15000 Hz and the second pulse peaked slightly higher (Fig. 2).

**Table 2.** Numbers of sounds produced by *Neolamprologus pulcher* associated with behaviours in the laboratory experiment

| Behaviour | #1_M | #2_F | #3_M | #9_F | #10_M | Total |
| --- | --- | --- | --- | --- | --- | --- |
| Soft touch | 0 | 0 | 0 | 1 | 0 | 1 |
| Parallel swim | 0 | 0 | 0 | 1 | 0 | 1 |
| Puffed throat | 0 | 0 | 2 | 0 | 8 | 10 |
| Aggressive posture | 0 | 8 | 5 | 0 | 20 | 33 |
| Lateral display | 2 | 6 | 15 | 1 | 9 | 33 |
| Pseudo-mouth display | 1 | 0 | 0 | 0 | 1 | 2 |
| Submissive posture | 0 | 6 | 0 | 3 | 0 | 9 |
| Total pulses with behaviour | 2 | 20 | 20 | 6 | 34 | 82 |

At times multiple behaviours were displayed conjointly with a given pulse, for example both a lateral display and a puffed throat. M = male and F = female

**Table 3.** Parameters (mean ± SD) of the acoustic signals associated with and without a typical *Neolamprologus pulcher* social behaviour

|  | No.<br>fish | Total<br>pulses | Pulse<br>duration<br>[ms] | Pulse peak<br>frequency<br>[Hz] | Total<br>calls | Call<br>duration<br>[ms] | Pulses<br>per call |
| --- | --- | --- | --- | --- | --- | --- | --- |
| With<br>behaviour | 5 | 82 | 11.5 ± 3.5 | 12280.5 ±<br>3740.3 | 43 | 896.0 ±<br>804.4 | 2.0 ±<br>0.7 |
| Without<br>behaviour | 2 | 10 | 13.2 ± 2.8 | 13992.2 ±<br>1889.3 | 5 | 294.4 ±<br>324.0 | 2.0 ±<br>0.0 |
| Pooled | 6 | 92 | 12.0 ± 3.4 | 12938.7 ±<br>3494.0 | 48 | 836.0 ±<br>733.7 | 2.0 ±<br>0.7 |

One fish emitted sound both with and without behaviour, therefore the pulses for each were calculated separately, except when pooled

**Fig. 3.**
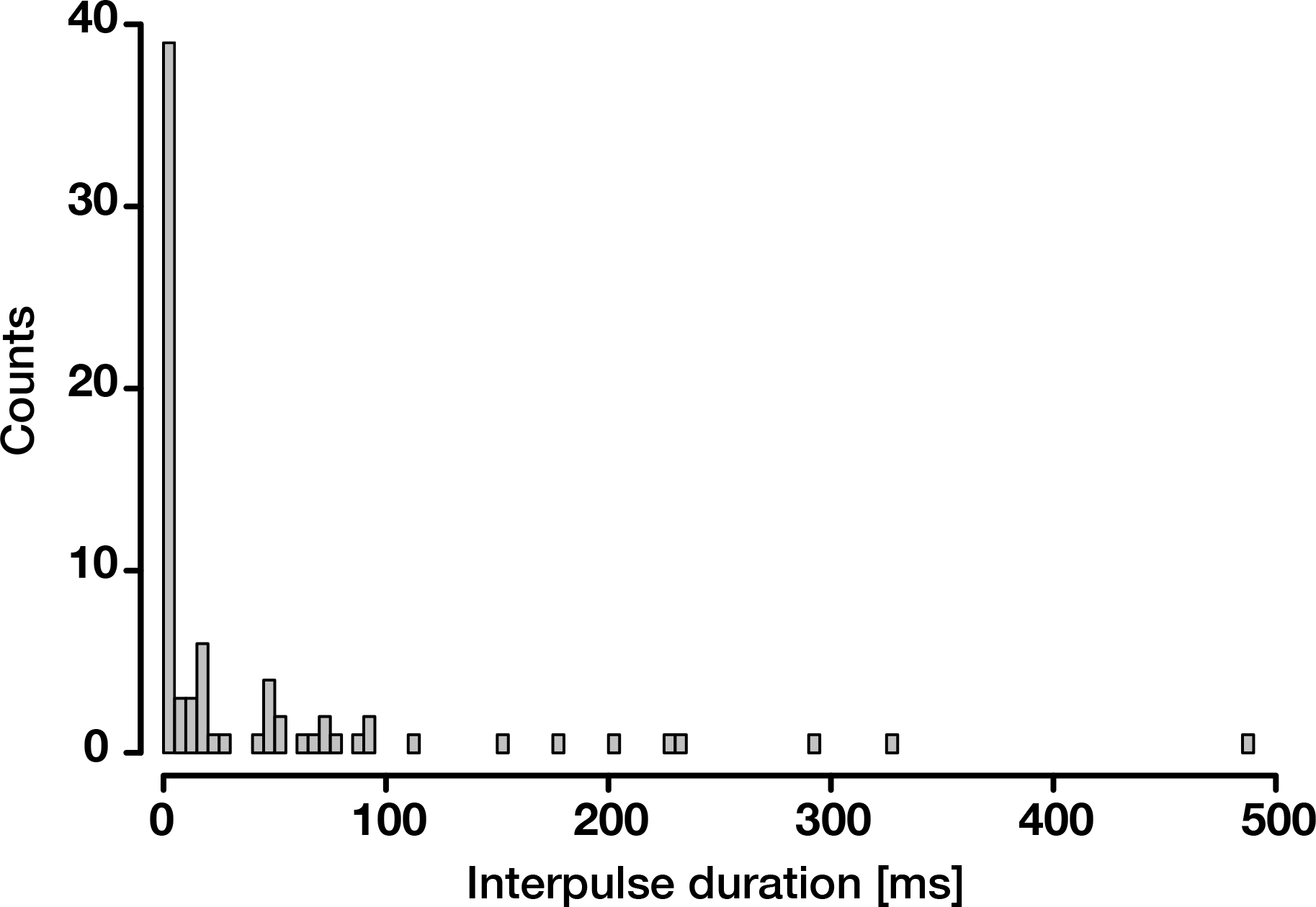
Histogram of interpulse duration frequency. The majority of pulses within a call are shortly separated by less than 0.4 s

Male *N. pulcher* produced more and longer pulses than females, however the peak frequencies of the pulses were very similar in both sexes (Table 4). The standard two-pulsed call was found in both sexes, although males had more calls than females, as well as a longer call duration (Table 4).

**Table 4.** Sex differences in the parameters (mean ± SD) of the acoustic signals of *Neolamprologus pulcher* in the laboratory experiments

|  | No.<br>fish | Total<br>pulses | Pulse<br>duration<br>[ms] | Pulse peak<br>frequency<br>[Hz] | Total<br>calls | Call<br>duration<br>[ms] | Pulses<br>per call |
| --- | --- | --- | --- | --- | --- | --- | --- |
| Male | 4 | 64 | 14.1 ± 2.1 | 12710.0 ±<br>4303.6 | 36 | 918.0 ±<br>770.4 | 1.8 ±<br>0.1 |
| Female | 2 | 28 | 8.5 ± 1.5 | 13396.2 ±<br>2201.8 | 12 | 669.6 ±<br>910.2 | 2.3 ±<br>1.4 |
| Pooled | 6 | 92 | 12.0 ± 3.4 | 12938.7 ±<br>3494.0 | 48 | 836.0 ±<br>733.7 | 2.0 ±<br>0.7 |

All sounds produced were taken into account, both with and without a typical social behaviour

## Discussion

### Sound production by Princess cichlids

Multimodal communication is expected in socially complex species (Freeberg et al., 2012) that live in unstable environments (Munoz & Blumstein, 2012). In this study we report the production of sounds often associated with a visual display by the cooperatively breeding Princess cichlid, *N. pulcher.* Our analyses confirm recent findings that this species does not produce the low frequency sounds common to many other cichlids or fish species in general (Pisanski et al., 2014), for which we suggest the term “low frequency silencing”. However, we found strong evidence for deliberate production of high frequency double-pulse calls by *N. pulcher.* In our field and laboratory experiments we found that both males and females produce high frequency sounds (above 5 kHz, average ~12 kHz) in an agonistic context induced by mirrors.

High frequency sound production has long been reported in cichlids, including in species from Lake Tanganyika (e.g. (Myrberg, Jr. et al., 1965; Nelissen, 1978)). Peak frequencies are similarly high (above 5 kHz, often higher than 20 kHz) but temporal characteristics differ substantially among species. *Neolamprologus pulcher* produces a distinct double-pulse clicking call while others *(Astatotilapia burtoni, Simochromis diagramma,* different *Tropheus* spp.) produce a creaking or chewing multi-pulsed call (Nelissen, 1978). These short pulses of sound and high frequency in *N. pulcher* point towards a stridulatory mechanism of sound production. It has been suggested that African cichlids may produce sound by rubbing together the teeth on their pharyngeal jaws (Rice & Lobel, 2004), although this mechanism is yet to be confirmed. (Fine & Parmentier, 2015) suggest that stridulatory mechanisms should contain a wide range of frequencies, such as the broadband sound produced by *N. pulcher.*

Most of the sounds recorded in this study were produced in association with an aggressive visual display, but interestingly also in submissive displays. Importantly, since fish also produced sound with similar characteristics without an associated behaviour, we can infer that sound production is not a sole by-product of a visual display but instead can be generated independently. By examining both wild and captive fish we could also exclude any effect of captivity and captive breeding on “low frequency silencing” in *N. pulcher.* The evolutionary reasons for loss of low frequency sounds are still unclear.

### Acoustic differences between and within wild and captive individuals

Both wild and captive individuals generate characteristic high frequency double-pulse clicks, but pulses of *N. pulcher* in the laboratory recordings were longer in duration compared to the field recordings (one order of magnitude on average). Interestingly, male and female *N. pulcher* differed also in temporal parameters. Cichlid acoustic studies have shown variation in pulse duration between closely related species, suggesting it is evolutionarily labile: mean pulse duration in *Oreochromis mossambicus* is 150 ms, compared to 10ms in *Oreochromis niloticus* (Amorim et al., 2003; Longrie et al., 2008), and species in the genus *Maylandia* show 2-3 times differences in mean pulse duration (Danley et al., 2012). Furthermore, context-and sex-specific differences have been reported in *Maylandia (Pseudotropheus) zebra* (Simões et al., 2008), and intra-individual variation in sound duration and pulse rate in response to motivation has been demonstrated in three distantly related cichlid species (Myrberg, Jr. et al., 1965). It is thus possible that noisier captive conditions have induced changes on labile temporal properties of *N. pulcher* sounds (pulse duration/period) in a similar way that environmental noise has impacted call duration and rate in Cope’s grey treefrog, *Hyla chrysoscelis* (Love & Bee, 2010) or song amplitude in common blackbird, *Turdus merula* and other birds (Nemeth et al., 2013).

### Significance of high pitch sounds

Reports of low (i.e. below 2-3 kHz) frequency sounds in cichlid fishes have been dominating the literature in recent years. This has likely both technical and biological explanations. On one hand, it is possible that sounds produced by cichlids in a reproductive context are mostly low frequency (e.g. (Nelissen, 1978)), while recording of narrower bandwidths or applying low-pass filters to raw data could account for masking of higher frequencies (Ripley & Lobel, 2004; Amorim et al., 2008; Longrie et al., 2008, 2009; Simões et al., 2008; Bertucci et al., 2012; Maruska et al., 2012; Pisanski et al., 2014). But perhaps the overarching reason relates to the expectation that fish are sensitive only to low frequency sounds and cannot hear above a certain threshold (e.g. (Heffner & Heffner, 1998)), which would render such high frequency sounds biologically irrelevant. It is presently unclear whether *N. pulcher* can detect such high frequencies, as hearing sensitivities have not been studied in this species and those of the close-relative *N. brichardi* (Gante et al., (n.d.)) have been investigated only in the range 100-2000 Hz (Ladich & Wysocki, 2003). Nevertheless, evidence has been mounting that some species react to high frequency sounds: for instance, behavioural studies indicate that cod *Gadus morhua* can detect ultrasonic signals up to 39 kHz and the clupeid *Alosa sapidissima* of over 180 kHz, well past human hearing (reviewed in (Popper & Lu, 2000)). Furthermore, new data indicate that species might have multiple hearing maxima, as bimodal w-shaped sensitivity curves have been described in Malawian cichlids previously thought to have only a u-shaped sensitivity curve peaking at low frequencies (van Staaden et al., 2012).

Nelissen (Nelissen, 1978) suggested that vocal complexity (measured as number of sound types) in six cichlid species from Lake Tanganyika varies inversely with number of colour patterns, such that different species would specialise along one of the two communication axes. Maruska et al. (Maruska et al., 2012) showed that acoustic signalling is an important sensory channel in multimodal courtship in the cichlid *A. burtoni.* Females responded to sounds even before seeing males (Maruska et al., 2012), which suggests that sounds could function as a long-distance attraction signal in the turbid waters of river deltas inhabited by this species. Sounds in the cooperative breeding *N. pulcher* could play a role in multimodal communication in an agonistic context and to maintain group cohesion. Since *N. pulcher* also produced sound in the confines of the shelter, it is possible that individuals can use acoustic signals when retreating to their shelter and other forms of communication are limited. Importantly, high frequency signals would also transmit more efficiently above the low frequency background noise of the underwater world, particularly in windy conditions (van Staaden et al., 2012) or crowded fish neighbourhoods. These longer-range high pitch sounds would allow communication among individuals belonging to different family groups, establishing a chorus across the colony.

While the ability of *N. pulcher* to hear in this high frequency range is still to be determined, several hearing ‘specialists’ inhabiting Lake Tanganyika could be potential interspecific receivers of the acoustic signals generated by cichlids. Hearing specialists that can detect sounds in the kHz generally have their swim bladder acoustically coupled to the inner ear (Popper & Lu, 2000). These include several catfish of the families Malapteruridae, Mochokidae, Claroteidae and Clariidae (Siluriformes) that can hear higher frequency sounds and predate on cichlids. Other potential candidates would be the many species that lurk around *Neolamprologus* rocky habitat, such as spiny eels of the family Mastacembelidae (Synbranchiformes) and perches of the family Latidae (Perciformes).

## Conclusion

We have shown that *N. pulcher* produces high frequency (above 5 kHz, average ~12 kHz) double-pulsed calls. Sounds are most often produced jointly with aggressive or submissive visual displays, although both acoustic and visual signals can be produced in isolation. It is unclear whether the receiver of such sounds is intra-or interspecific given our general lack of understanding of hearing sensitivities of fishes inhabiting Lake Tanganyika. In the event that cichlids can hear such high pitch sounds, an as of yet undescribed morphological adaptation is expected to exist. Non-visual sensory modalities in African cichlids may thus have a larger impact than originally expected and could be an important aspect in their adaptive radiation.

## Acknowledgements

We thank Lia Albergati and Benjamin Küng for assistance in the field and Miguel Vences for use of the recording equipment. This project was funded by a travel grant from the University of Basel to RKS, by grants from the European Research Council (ERC; StG “CICHLID~X”) and the Swiss National Science Foundation (SNF) to WS, and by the “University of Basel Excellence Scholarships for Young Researchers” and “Novartis Excellence Scholarships for Life Sciences” to HFG.

## Additional files

**Additional file 1: Audio file.** Two double-pulsed calls of a male *Neolamprologus pulcher,* produced during an aggressive lateral display in the laboratory experiments. The first double-pulsed call corresponds to Fig. 2

**Additional file 2: Audio file.** One double-pulsed call of a female *Neolamprologus pulcher,* produced during submissive posture in the lab experiments.

**Additional file 3: Audio file.** One double-pulsed call of a male *Neolamprologus pulcher,* produced without behavioural display (motionless) in the lab experiments.

